# Sentimental Tweets Classification of Symptomatic COVID-19

**DOI:** 10.1101/2021.12.15.472745

**Authors:** P Tharun

## Abstract

The approach I described is straightforward, related to COVID-19 SARS based tweets and the symptoms, that people tweet about. Also, social media mining for health application reports was shared in many different tasks of 2021. The motto at the back of this observe is to analyses tweets of COVID-19 based symptoms. By performing BERT model and text classification with XLNET with which uses to classify text and purpose of the texts (i.e.) tweets. So that I can get a deep understanding of the texts. When developing the system, I used two models the XLNet and DistilBERT for the text sorting task, but the outcome was XLNET out-performs the given approach to the best accuracy achieved. Now I discover a whole lot vital for as it should be categorizing tweets as encompassing self-said COVID-19 indications. Whether or not a tweets associated with COVID-19 is a non-public report or an information point out to the virus. Which gives test accuracy to an F1 score of 96%.

## 1. INTRODUCTION

The COVID-19 pandemic has encouraged an astounding damage of anthropological life universal and grants an unexpected food system, worldwide health, and the workplace are all under threat [1]. As this disease is highly contagious and rapidly changing, one of the best sources for the live information is on the social media [2,4]. There can be numerous sources of symptoms information on social media such as news and scientific articles (facts), other people’s account (second or third person statements) and self-report (first person statements). In this paper we will discuss our method as a team contributing in SMM4H [5] shared task 6 related to the sorting of such information from social media platform like twitter. We will be looking at using pre-trained NLU models like BERT and XLNET for this task.

So far, the world as we know it has changed, and today we live in a new and ever-changing atmosphere. The way we live, communicate and talk to others has changed forever [4,5]. In these special circumstances, virus risk plays a role in educating, displaying, and refusing to distribute data to the public. The coronavirus poses a real pandemic risk and poses a major challenge to the government to control, prepare, respond and improve. Health groups, stakeholders and the media [5].

The current catastrophe is due to COVID-19, the health community is being exposed to socially unconventional circumstances. Almost everything in our daily lives is becoming globalized, and people are relocating and travelling more often, Globalization means the interconnectivity between economies is growing, and tool development and knowledge are becoming more sophisticated.

From the sort of predicted class label, it is clear that in addition to semantics, contextual illustration also plays an imperative role. This task usually uses repetitive patterns, which are calculated along through the character positions of the input and output sequences. In the intention time step, they produce a classification of concealed circumstances that based on the input of the previous hidden state ht-1 and position t. This inherent chronological nature prevents the parallelization of training samples, which becomes decisive in longer classifications. Because memory limitations limit the group processing in the example. Initially, there are two strategies for contextual representation to apply previously trained language representations to future tasks: function-based and fine-tuning. However, both are limited by the fact that they are one-way language models and cannot learn universal language representations. In Recent developments in generalized autoregressive model (XLNET) views resolve these two problems since it captures bi-directional framework by means of a mechanism called “permutation language modeling” Determine the depth of unlabeled text by adjusting the left and right context together at all levels [6]. And encode to attain the vector illustration of the verdict, and fix the BERT for the ternary classification problem.

Similarly, due to the outbreak of this virus, false reports and criticisms of movements that disrupt communication between health authorities and cause social tension continue to appear [7]. Through the sentiment analysis of the new virus, five main problems have been identified, Positive to negative [8]. An extensive analysis of tweets from Indian users of social media was conducted in this article. To utilize deep learning for Text Data Analysis, we defined a generalized autoregressive model enhanced by transformer-XL model pre-trained (XLNET). The performance of numerous various Machine Learning algorithms is compared with BERT by using logistic regression, vector machine support, and single-layer long-term memory algorithms. Therefore, it is hoped that the following questions will be answered in this article.

These models are trained and retrained by XLNet to perform and come with best accuracy to achieved both from positive and negative to attain the accuracy level.

- Q1: What remain the most prevalent keywords in global tweets?
- Q2: What are the ramifications of these tweets on public health?
- Q3: Using machine learning algorithms, can emotions, feelings, and thoughts be analyzed?
- Q4: Are deep learning BERT models as effective as the other three traditional machine learning models?

### Fake News in Social Media Correlated with COVID-19 Studies

The rise of false information through Internet sites is becoming a global problem. Although fake news is not essential, it is currently worrying because online media is known for cooperating and spreading new ideas. [9] The COVID19 pandemic shows the valuable impact of the new data. Dissemination of false information will obviously weaken personal awareness and undermine government countermeasures are feasible. When misinformation on social media spreads, panic and fear can be caused among COVID19 patients, alerting authorities and prompting citizens to confirm the spread. [10] With the COVID19 pandemic [11–14]. The spread of information about COVID19 through immense data analysis on large social media platforms can be personalized on a large scale to investigate rumors about the epidemic [15].

It is a well-known social media platform and Weibo system anywhere people can post and exchange communications called “tweets”. Twitter receives 500 million tweets a day and 200 billion tweets a year due to how it is structured turn out to be the main data access point for online media discussions with the public and global environments [16]. Unfortunately, due to the blowout of fake news, it is also a major source of global panic. It is described in 2020 that most tweets about COVID19 are in a positive mood, but users often contribute in the spread of negative tweets, and when manipulative the frequency of words in the tweets, they find that there are not many useful words. [17]

As with any virus, the current pandemic has spawned rapidly spreading rumors and conspiracy theories. My research has been centered on automatic detection and categorization of diseases and viruses using machine learning methods [16,17] and identify the narrative framework that supports this false propaganda. The consequences of misleading information about COVID19 and its dogmatic philosophy eventually festering community well-being. [18]

In 2020 WHO reported that there are a lot of rumors and false stories about COVID19 circulating on social media. It is problematic to distinguish this fake news from real news, but its accuracy or reliability is no longer possible. [19] Counterfeiting on societal mass media sites can benefit individuals, health professionals, and administrations avoid unnecessary psychological stress. These programs provide general explanations for any non-exclusive events and can be easily customized based on conspiracy theories. In the works verified by tweets, I tested the deep learning type of machine learning model and verified the effectiveness of the model by comparing it with three other traditional models.

## 2. Task and Data Description

### A. Task

Tasks of the Social Media Mining for Health Applications (SMM4H) 2021 overall tasks. [20] requires participants to develop an automated classification system to identify mentions of

1. the self-report,
2. Non-personal reports, as well
3. Bibliography. COVID-19 SARS news, articles, and related topics are mentioned.

It is expressed by way of a multi-class sorting problem, in which the system must predict the label of each tweet in order to set up tweets.

In this study, I included the tweets data of twitter users during the COVID19 outage in the nation. The dataset of 4,010 tweets is taken and contains clean tweets on issues such by way of COVID19, Coronavirus, blocking, etc. For the analysis, I considered the tweets from the Kaggle platform Twitter to record the coronavirus and conclusions.

I used natural learning processing technology (NLP), which is a form of machine learning that helps process tweets. Generally, NLP includes various text mining methods, such as clatter reduction, halt words, and buzzword elimination. In direction to recover the recital of the BERT model, measurements, intangible factors, or text noise such as accents, math, and line breaks have been removed. [21,22] Deleting these items will reduce the testing area for potential skills, thereby improving performance.

I divided the record of each task into each part of approximately 1500 and 1780 annotated examples, set up models for the 4 convolutions, and checked the remaining convolutions. For each fold, I optimize the model in more and more training examples, in addition, for these tasks, [23,24] I tried to use the previously trained model for another task that I suspect may be useful, because these tasks seem to be related.

As mentioned earlier, in order to usage pre-trained text weights in the BERT model, we must adjust the input attributes of the encoding. I applied 600 tokens to the maximum length on this particular [25] model. In addition, I analyzed the token length of each tweet. A large number of tweets contain less than 600 tokens.

**Fig 2.1:**
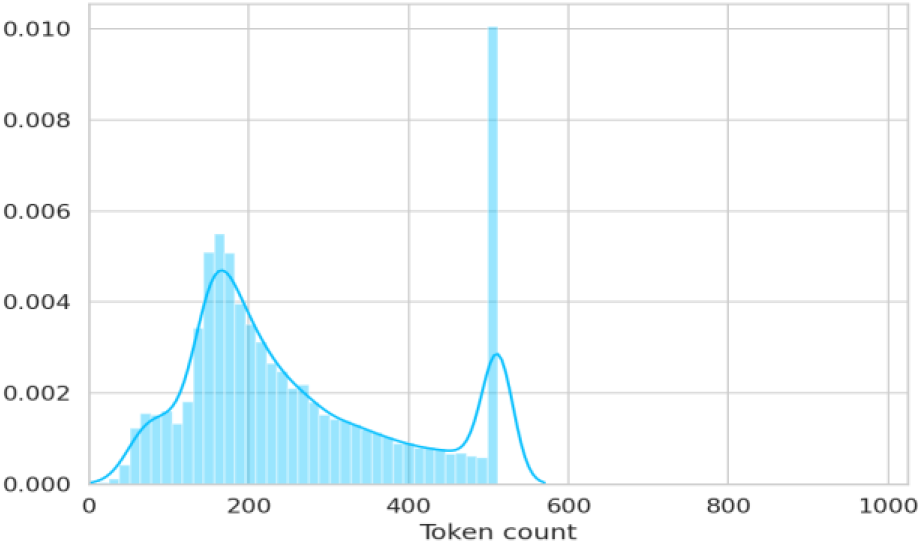
The result of the total length of tweets

Based on the given multi-class classification problem, I used a dependent variable whose values were positive, extremely positive, neutral, negative, and extremely negative. In addition, I divided this question into two binary classifications: positive and negative. Towards improving accuracy, the classification is further separated into three categories: positive, negative, and neutral. The results of these algorithms will be compared in the evaluation phase.

To clean up, I deleted some of the sentences from “@” to the first space, links, numbers, “#” and extra spaces. Later in the preprocessing, I converted the data into a machine-readable format. I transformed the label to three discrete integers. And I use the tokenizer described in the model section to tweet the text in the token. Then I cropped the tweet text to reduce indentation. For BERT, I trimmed before applying the tokenizer and provide a proposal of length 66, while for XLNet, I trimmed after using the tokenizer to reach a sequence length of 160.

### B. Data Description

The training data set contains 9,000 labeled tweets, the validation data set contains 5.76 labeled tweets, and the test data set contains 6,500 unlabeled tweets.

The classification model cannot be adapted to the data set for confirmation. I divided the data set into two parts: working out and challenging. Well-trained tweets help classify data patterns, reduce errors, and test data sets for analysis. 85% of tweets are used for educational purposes and 17% are used for testing determinations. To Provide demographic data by tweet multi-class classification [26,27].

### C. BERT Model

My first system for this task is BERT. BERT stands for Bidirectional Transformer Encoder Illustrations, which is trained by using the left and right contexts of the masked words to randomly predict the masking tags during pre-training. [28,29] Token also has a second purpose-to predict whether two given sentences are continuous. In my experiments, I used uppercase and lowercase letters in this model. The basic version of BERT has 12 encoder levels, 768 hidden level measurements, and 12 attention levels. The head has 109 million parameters, while the big head has 24 encoder levels, 1024 hidden layer measurements and [30]16 attention heads, with 335 million parameters. I use their tokenizer.

### D. XLNet Model

XLNet is an extension of the previously trained Transformer XL model that uses autoregression to learn bidirectional framework by exploiting the anticipated probability of entirely variations of the input sequence decomposition order.

It allows to use autoregressive and automatic pre-workout coding techniques to overcome inconsistencies in pre-workout fine-tuning.

This can be easily used for any task by loading a pre-trained model and setting it as a future task. In short, hugging face Transformers has provided several model classes to perform specific tasks for subsequent use of XLNet. I only need to download and configure them without writing our own model classes, which are additional layers on top of the XLNet model.

I used XLNet as the second system. XLNet is an autoregressive model, which is different from BERT in that it uses an autoregressive formula to learn two-way context. The output of the word tag is calculated captivating into interpretation the arrangement of all word tags in the sentence, which is contrary to the traditional method that is only used on the left or right side of the target tag. We experimented with the basic and large versions of this model. The basic version has 12 layers, 778 hidden layer sizes and 12 consideration heads, 110 million. We use their tokenizer.

## 3. Model Training

We did some experiments on clean and unclean data. We checked the basic and major versions of BERT and XLNet, as well as numerous training methods, fine-tuning and retraining of the complete prototypical. The outcomes are made known in Table 1. Use the appropriate sorting level. For all these experiments, the loss function represents the loss of cross entropy, and the learning rate of the optimizer. I found that the larger version of BERT retrained with the original data performed better. The XLNet version of the tokenizer with the basic version of the model works best. Thus obtained the best results by retraining the model using the original data. The best system results of the test data for general problems are shown in Table 2.

**Fig 3.1:**
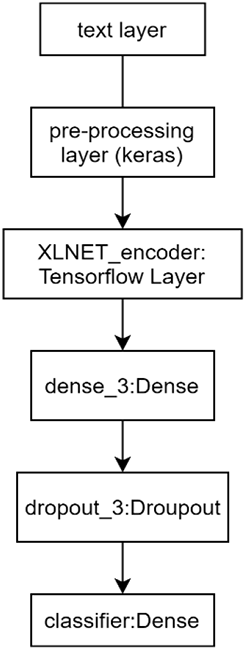
Model

**Table 1:** This gives the statistics of the dataset.

| Dataset | LN | NP | Self | Total |
| --- | --- | --- | --- | --- |
| Train | 4267 | 3404 | 1248 | 8919 |
| Validation | 250 | 177 | 88 | 515 |
| Test | - | - | - | 6500 |

**Table 2:**
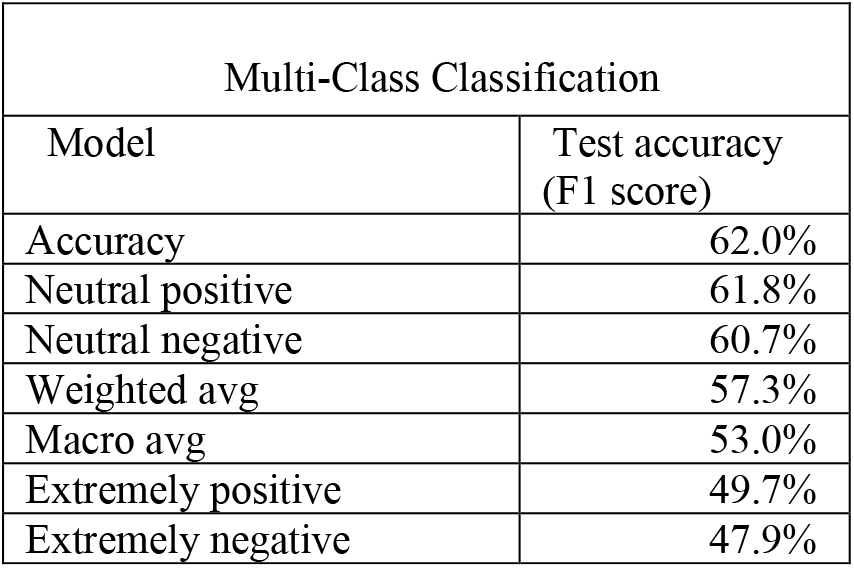
Demographic representation of tweets multi-class classification.

| Multi-Class Classification |  |
| --- | --- |
| Model | Test accuracy (F1 score) |
| Accuracy | 62.0% |
| Neutral positive | 61.8% |
| Neutral negative | 60.7% |
| Weighted avg | 57.3% |
| Macro avg | 53.0% |
| Extremely positive | 49.7% |
| Extremely negative | 47.9% |

A model of BERT identifies whether the second set of a pair of operators in the source file corresponds to the final result after collecting a pair of operators as input. During training, partial of the information sources are paired so that the reverse sentence is the result judgement in the first file, and the other half selects the incorrect sentence from the amount as the reverse sentence. Consider separating the incorrect sentence from the main sentence. BERT’s input source is a combination of inserted markers, segments, and positions. During training, the training data is processed as follows so that the model can distinguish between different topics.

Position insertion is added to each mark to indicate its status consistently. The BERT tokenizer performs word segmentation on the part of the word: the dictionary displays discrete language features, and again adds high-frequency word combinations. Use word cloud graphs to rank keywords in tweets. For training and verification purposes, there are two subsets of data. Use the accuracy calculation to see how accurate the model is. Retraining is also superior to refining according to my observations. BERT, the large version is better than the basic version, and on XLNet, the basic version is better than the large version. The test suite, and XLNet does a better job in the test suite.

## 4. SYSTEM OVERVIEW AND IMPLEMENTATIONS

### A. System Overview and Design

We studied the automatic coding and autoregressive models of classification problems, and selected BERT and XLNet. In our experiments, we used the previously trained hugging face model [30]. They use uppercase, case-sensitive versions. 103 This is necessary because capital letters are important for defining nouns and pronouns, which in turn are important for defining the first, second, and third persons in the sentence.

I also studied issues related to community sentimentality, shimmering deep concerns about the coronavirus and COVID19, foremost to increased anxiety and negative emotions. I also established the use of exploratory and eloquent text analysis and techniques. Visualize text data to discover early ideas. Group words by the level of specific non-text variables. Using those datasets to train to the best accuracy achieved using XLNet also outperforms the other model.

Many potential ethical issues have been identified regarding how professionals use Twitter data for research; these include the use of tweets by vulnerable groups in crisis situations [31,32]. Research, as well as human beings, are no longer the responsibility of researchers, and researchers are no longer the responsibility of “data subjects” [33]. As a result, public data is valuable as a voluntary contribution, even though it does not violate ethical principles. From Twitter users, access public spaces. Moreover, the study shows how Twitter data was analyzed in connection with pandemics, including the 2009 swine flu [34,35], demonstrating a mature mentality toward identifying and managing infections and crises by using social media.

The first layer and the second layer both combined to form a XLNet based framework by pre-processing to analyze the correct sequence model to the given based framework. Which utilizes the second layer to filter by using keras layer to sort out the dense and classifying the tweets by another encoder to predict what come next as positive or negative based on the situation it then again TensorFlow for sorting out the tweets again by repetition method this goes until by identifying the accuracy by training iteratively.

**Figure 4.1:**
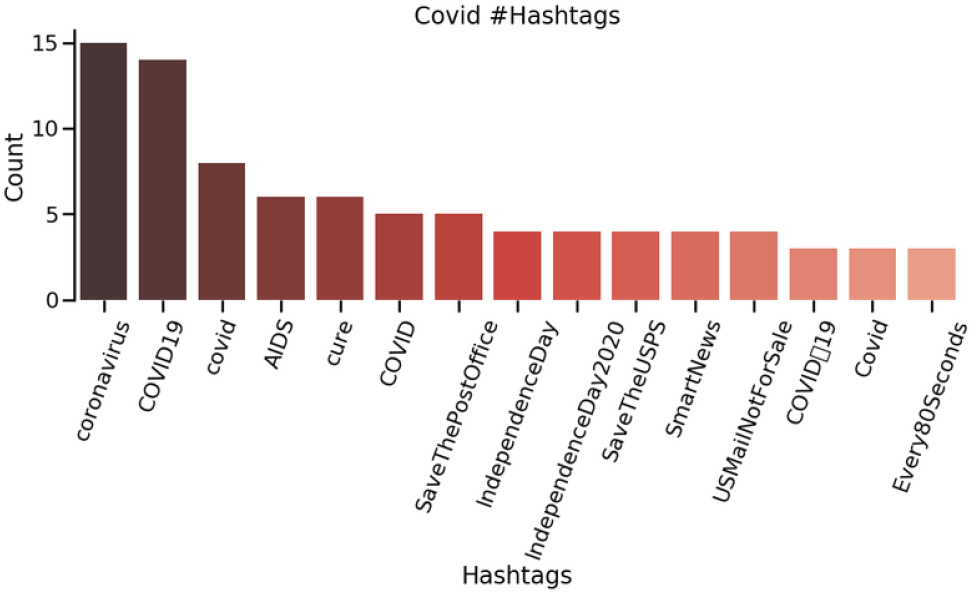
This depicts COVID-19 is mainly used as a hashtag in tweets, people also want to talk about other events in tweets.

In addition to, BERT base, XLnet and Electra, we gained control of them all. FP16 calculations are used to reduce the size [36] of the model as controlled fine-tuning consumes a lot of GPU memory. To train at 0.00003, we selected the 0.00003 settings. Sequences can have a maximum length. 250 is the set value. We have set the number of sublots to two. The learning rate is set to 3e5 using Adam optimizer. Training is done over three epochs with gradient accumulation.

## 7. CONCLUSION

So, as discussed, a simple way to adapt the XLNet model to 2021 social media mining common problems in healthcare applications. These results are not up-to-date, but they are competitive and demonstrate the benefits of using contextualized and pre-trained language models. On a large scale. The model has achieved F1 score of 93% accuracy (as shown in table 1). I also discussed the benefits of first training the model for the task at hand and determine when it might be useful.

**Table 1:** Results

| Dataset | BERT | XLNet |
| --- | --- | --- |
| Result | .941 | .968 |
| Test | .666 | .784 |

## REFERENCES

1. Chawla, S.; Mittal, M.; Chawla, M.; Goyal, L. Corona Virus-SARS-CoV-2: An Insight to Another way of Natural Disaster. EAI Endorsed Trans. Pervasive Health Technol. 2020.

2. Yuan C, Wu Y, Qin X, Qiao S, Pan Y, Huang P, Liu D, Han N (2019) An effective image classification method for shallow densely connected convolution networks through squeezing and splitting techniques. Appl Intell 49(10):3570–3586.

3. Lin E, Chen Q, Qi X (2020) Deep reinforcement learning for imbalanced classification. Appl Intell, 1–15.

4. Staszkiewicz, P.; Chomiak-Orsa, I. Dynamics of the COVID-19 Contagion and Mortality: Country Factors, social media, and Market Response Evidence from a Global Panel Analysis. IEEE Access 2020, 8, 106009–106022.

5. Guo, Y.-R.; Cao, Q.-D.; Hong, Z.-S.; Tan, Y.-Y.; Chen, S.-D.; Jin, H.-J.; Tan, K.-S.; Wang, D.-Y.; Yan, Y. The origin, transmission and clinical therapies on coronavirus disease 2019 (COVID-19) outbreak—An update on the status. Mil. Med. Res. 2020, 7, 1–10.

6. Mittal, M.; Battineni, G.; Goyal, L.M.; Chhetri, B.; Oberoi, S.V.; Chintalapudi, N.; Amenta, F. Cloud-based framework to mitigate the impact of COVID-19 on seafarers’ mental health. Int. Marit. Health 2020, 71, 213–214.

7. Liu P, Zhang H, Eom KB (2016) Active deep learning for classification of hyperspectral images. IEEE J Sel Top Appl Earth Obs Remote Sens 10(2):712–724.

8. Akande, O.N.; Badmus, T.A.; Akindele, A.T.; Arulogun, O.T. Dataset to support the adoption of social media and emerging technologies for students’ continuous engagement. Data Brief 2020, 31, 105926

9. Gorriz M, Carlier A, Faure E, Giro-i-Nieto X (2017) Cost-effective active learning for melanoma segmentation. arXiv:1711.091681711.09168.

10. Huong Dang, Kahyun Lee, Sam Henry, and Özlem Uzuner. 2020. Ensemble BERT for classifying medication-mentioning tweets. In Proceedings of the Fifth Social Media Mining for Health Applications Workshop & Shared Task, pages 37–41, Barcelona, Spain (Online). Association for Computational Linguistics.

11. Jacob Devlin, Ming-Wei Chang, Kenton Lee, and Kristina Toutanova. 2019. BERT: Pre-training of 140 deep bidirectional transformers for language understanding. In Proceedings of the 2019 Conference of the North American Chapter of the Association for Computational Linguistics: Human Language Technologies, Volume 1 (Long and Short Papers), pages 4171–4186, Minneapolis, Minnesota. Association for Computational Linguistics.

12. Arjun Magge, Ari Klein, Ivan Flores, Ilseyar Alimova, Mohammed Ali Al-garadi, Antonio MirandaEscalada, Zulfat Miftahutdinov, Eulàlia FarréMaduell, Salvador Lima López, Juan M Banda, Karen O’Connor, Abeed Sarker, Elena Tutubalina, Martin Krallinger, Davy Weissenbacher, and Graciela Gonzalez-Hernandez. 2021. Overview of the sixth social media mining for health applications (# smm4h) shared tasks at naacl 2021. In Proceedings of the Sixth Social Media Mining for Health Applications Workshop & Shared Task.

13. Henry Tsai, Jason Riesa, Melvin Johnson, Naveen Arivazhagan, Xin Li, and Amelia Archer. 2019. Small and practical bert models for sequence labeling. arXiv preprint arXiv:1909.00100.

14. Sebastian Ruder. 2021. Recent Advances in Language Model Fine-tuning.

15. Victor Sanh, Lysandre Debut, Julien Chaumond, and Thomas Wolf. 2019. Distilbert, a distilled version of bert: smaller, faster, cheaper and lighter. arXiv preprint arXiv:1910.01108.

16. Kuo W, Häne C, Yuh E, Mukherjee P, Malik J (2018) Cost-sensitive active learning for intracranial hemorrhage detection. In: International conference on medical image computing and computer-assisted intervention. Springer, New York, pp 715–723.

17. Lv X, Duan F, Jiang JJ, Fu X, Gan L (2020) Deep active learning for surface defect detection. Sensors 20(6):1650.

18. Yoo D, Kweon IS (2019) Learning loss for active learning. In: Proceedings of the IEEE conference on computer vision and pattern recognition, pp 93–102.

19. Zhan Shi, Xinchi Chen, Xipeng Qiu, and Xuanjing Huang. 2018. Toward diverse text generation with inverse reinforcement learning. IJCAI

20. Zhao Z, Yang X, Veeravalli B, Zeng Z (2020) Deeply supervised active learning for finger bones segmentation. arXiv:2005.03225.

21. Roman Novak, Michael Auli, and David Grangier. 2016. Iterative refinement for machine translation.

22. Ratish Puduppully, Li Dong, and Mirella Lapata. 2019. Data-to-text generation with content selection and planning. In AAAI, volume 33, pages 6908–6915.

23. Garcia, L.P.; Duarte, E. Infodemic: Excess quantity to the detriment of quality of information about COVID-19. Epidemiol. Serv. Health 2020, 29

24. Hung, M.; Lauren, E.; Hon, E.S.; Birmingham, W.C.; Xu, J.; Su, S.; Hon, S.D.; Park, J.; Dang, P.; Lipsky, M.S. Social Network Analysis of COVID-19 Sentiments: Application of Artificial Intelligence. J. Med. Internet Res. 2020, 22, e22590.

25. Di Domenico, G.; Sit, J.; Ishizaka, A.; Nunan, D. Fake news, social media and marketing: A systematic review. J. Bus. Res. 2021, 124, 329–341.

26. Apuke, O.D.; Omar, B. Fake news and COVID-19: Modelling the predictors of fake news sharing among social media users. Telemat. Inform. 2021, 56, 101475.

27. Zaman, A. COVID-19-Related Social Media Fake News in India. J. Media 2021, 2, 7.

28. Sohn K, Lee H, Yan X (2015) Learning structured output representation using deep conditional generative models. In: Advances in neural information processing systems, pp 3483–3491.

29. Depoux, A.; Martin, S.; Karafillakis, E.; Preet, R.; Wilder-Smith, A.; Larson, H. The pandemic of social media panic travels faster than the COVID-19 outbreak. J. Travel Med. 2020, 27, taaa031.

30. Gao, J.; Zheng, P.; Jia, Y.; Chen, H.; Mao, Y.; Chen, S.; Wang, Y.; Fu, H.; Dai, J. Mental health problems and social media exposure during COVID-19 outbreak. PLoS ONE 2020, 15, e0231924.

31. Ahmad, A.R.; Murad, H.R. The Impact of Social Media on Panic During the COVID-19 Pandemic in Iraqi Kurdistan: Online Questionnaire Study. J. Med. Internet Res. 2020, 22, e19556.

32. Cinelli, M.; Quattrociocchi, W.; Galeazzi, A.; Valensise, C.M.; Brugnoli, E.; Schmidt, A.L.; Zola, P.; Zollo, F.; Scala, A. The COVID-19 social media infodemic. Sci. Rep. 2020, 10, 1–10.

33. Twitter, Twitter Usage Statistics—Internet Live Stats. online:https://www.internetlivestats.com/twitter-statistics/ (accessed on 19 October 2020)

34. Chakraborty, K.; Bhatia, S.; Bhattacharyya, S.; Platos, J.; Bag, R.; Hassanien, A.E. Sentiment Analysis of COVID-19 tweets by Deep Learning Classifiers—A study to show how popularity is affecting accuracy in social media. Appl. Soft Comput. 2020, 97, 106754.

35. Shahsavari, S.; Holur, P.; Tangherlini, T.R.; Roychowdhury, V. Conspiracy in the time of corona: Automatic detection of COVID-19 conspiracy theories in social media and the news. J. Comput. Soc. Sci. 2020, 3, 279–317

36. Havey, N.F. Partisan public health: How does political ideology influence support for COVID-19 related misinformation? J. Comput. Soc. Sci. 2020, 3, 319–342

37. Obi-Ani, N.A.; Anikwenze, C.; Isiani, M.C. Social media and the COVID-19 pandemic: Observations from Nigeria. Cogent Arts Humanit. 2020, 7, 1799483.

38. Barkur, G.; Vibha; Kamath, G.B. Sentiment analysis of nationwide lockdown due to COVID 19 outbreak: Evidence from India. Asian J. Psychiatry 2020, 51, 102089.

39. Huynh, T.L. The COVID-19 risk perception: A survey on socioeconomics and media attention. Econ. Bull. 2020, 40, 758–764

40. Chintalapudi, N.; Battineni, G.; Di Canio, M.; Sagaro, G.G.; Amenta, F. Text mining with sentiment analysis on seafarers’ medical documents. Int. J. Inf. Manag. Data Insights 2021, 1, 100005

41. Devlin, J.; Chang, M.W.; Lee, K.; Toutanova, K. BERT: Pre-training of deep bidirectional transformers for language understanding. arXiv 2018, arXiv:1810.04805.

42. Chang, Y.W.; Hsieh, C.J.; Chang, K.W.; Ringgaard, M.; Lin, C.J. Training and testing low-degree polynomial data mappings via linear SVM. J. Mach. Learn. Res. 2010, 11, 1471–1490

43. Melamud, O.; Goldberger, J.; Dagan, I. Context2vec: Learning generic context embedding with bidirectional LSTM. In Proceedings of the 20th SIGNLL Conference on Computational Natural Language Learning 2016, Berlin, Germany, 11–12 August 2016.

44. India—COVID-19 Overview—Johns Hopkins. online: https://coronavirus.jhu.edu/region/india (accessed on 23 March 2021).

45. Samuel, J.; Ali, G.; Rahman, M.; Esawi, E.; Samuel, Y. COVID-19 Public Sentiment Insights and Machine Learning for Tweets Classification. Information 2020, 11, 314.

46. Abd-Alrazaq, A.; Alhuwail, D.; Househ, M.; Hamdi, M.; Shah, Z. Top Concerns of Tweeters During the COVID-19 Pandemic: Infoveillance Study. J. Med. Internet Res. 2020, 22, e19016.

47. Li, S.; Wang, Y.; Xue, J.; Zhao, N.; Zhu, T. The Impact of COVID-19 Epidemic Declaration on Psychological Consequences: A Study on Active Weibo Users. Int. J. Environ. Res. Public Health 2020, 17, 2032

